# Zika Virus Disease: A Simple Model Approach to Rapid Impact Assessment

**DOI:** 10.1101/044248

**Authors:** James M. Wilson, Robert W. Malone, Julie Smith-Gagen, Roman Pabayo, Zika Response Working Group

## Abstract

There has been substantial media attention regarding Zika infection as a threat to pregnancy, prompted by WHO’s declaration of a Public Health Event of International Concern (PHEIC). Here we present a simple risk assessment model for two states within the continental United States at-risk for autochthonous transmission of Zika virus, Texas and Florida. Our simple impact assessment model is partially validating at this early interval in the crisis.

## Introduction

Recent reports of high rates of primary microcephaly and Guillain-Barré syndrome associated with autochthonous transmission of Zika virus in French Polynesia and now in Central and South America have raised concerns that the virus circulating in these regions may represent a rapidly developing neuropathic, teratogenic, emerging public health threat.^1^ Brazil has represented a focus of interest within the broader of context of the crisis due to the intensity of Zika transmission and reported clinical outcomes. Brazil has reported 497,593 to 1,482,701 cases of Zika infection^2^, and from October 2015 to January 30, 2016, 5,640 cases of suspected perinatally acquired microencephaly^3^. Investigations for 1,533 of these cases resulted in exclusion of 950 cases (62%) due to over categorization of microencephaly or perinatal CNS malformation^3^, which has highlighted the challenges in determination of the true extent of perinatal impact of Zika infection. While investigations are ongoing to establish causality of Zika infection and microencephaly, the international community has adopted an assumption that Zika virus is the likely cause of the unusual increase in microencephaly and Guillain-Barré syndrome in Brazil and French Polynesia. ^3^ There are no licensed medical countermeasures available for Zika virus infection. ^1^ The purpose of this paper is to provide a preliminary impact assessment utilizing a simple mathematical model regarding expected importation of Zika virus for Texas and Florida.

The best-documented Zika virus animal reservoirs are primates, with predominant transmission to *via Aedes* species mosquito vectors.^4-8^ Forest-dwelling birds, horses, goats, cattle, ducks and bats have been demonstrated to be hosts of Zika virus, however it is unclear to what extend these hosts may serve as true ecological reservoirs. ^9^ Recent work indicates the potential for bloodborne^10^, sexual^11^, and oral^12^ transmission of Zika virus among humans. Perinatal transmission of the strain present in French Polynesia during the 2013-2014 outbreak has been documented, including the presence of Zika sequences in breast milk as demonstrated by PCR.^13^

While acknowledging research is still required to further elucidate possible additional ecological reservoir and vector species for Zika virus, we considered dengue and Chikungunya importation and autochthonous transmission in the United States to represent an appropriate initial reference point for an impact assessment. Dengue is endemic to tropical America and is the dominant arbovirus in the western hemisphere. Exotic Chikungunya is transmitting in the context of a “virgin soil” epidemic in tropical America^14^, including Brazil^15^. Dengue and Zika viruses are Flaviviruses; Chikungunya is a Togavirus. All three viruses are transmitted by *Aedes* species mosquitoes, with similar transmission dynamics.^16-18^

## Methodology

We used historical data from 2010-2015 for dengue and Chikungunya viruses in the United States^19^ to approximate impact of Zika transmission in Texas and Florida. This has been a period of unprecedented dengue importation, based on reported importations. From 2010 to 2015, these states have reported the highest absolute cases of importation and autochthonous transmission of dengue and importation of Chikungunya and therefore are proposed to be at highest risk of autochthonous transmission of Zika virus in the future. Autochthonous transmission of Chikungunya has not been documented in the United States to-date. We considered data for imported cases of dengue and Chikungunya, with a focus on the years of highest reported cases. Table 1 displays the data considered for this study, where the larger of the values (imported versus locally acquired dengue versus imported Chikungunya) was used to provide the most liberal estimates.

**Table 1.**
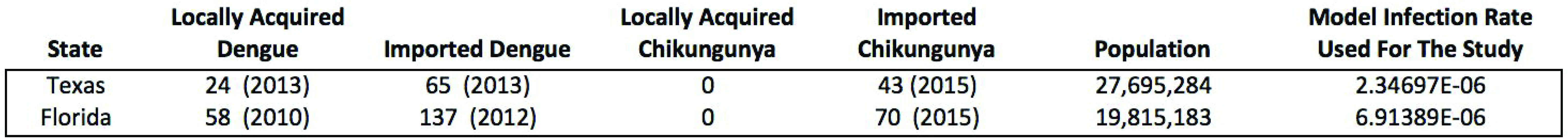
Data used for the simple risk assessment model. Imported dengue case counts were selected as the proxy for Zika virus importation for Texas and Florida.

Total state-level population and birth data were obtained from the respective state agencies.^20, 21^ Birth data was considered a proxy for pregnancies in each state and therefore, the population of interest for this analysis. To calculate these estimates, we multiplied the infection rate (as defined above) with the birth rate, followed by multiplication with the total state population (Table 2).

**Table 2.**
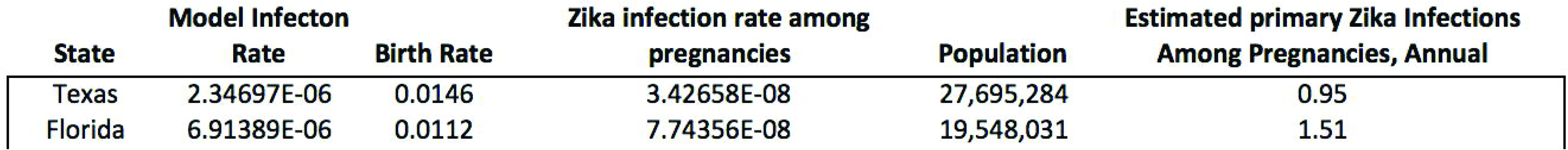
Simple model predictions based on maximized dengue and Chikungunya infection rates multiplied by the birth rates of Texas and Florida.

## Results

We estimate that annually, the cases of Zika infection among pregnancies will be low in Texas and Florida if Zika emergence follows a pattern similar to the United States’ experience thus far with dengue and Chikungunya (Table 1). The implication is we expect to observe no localized medical infrastructure strain in Texas or Florida due to importation of Zika virus disease and associated outcomes related to microencephaly or Guillain–Barré syndrome for the next three years. Based on observations of acute Zika community transmission abruptly ceasing following “virgin soil” outbreaks in French Polynesia^22^, we propose a similar pattern may be expected in Central and South America. It may be inferred from the experience in French Polynesia that this period of heightened transmission in the context of immunologically naïve populations in Central and South America will be limited to a period of several years.

## Discussion

As of February 24, 2016, there have been 107 confirmed imported cases of Zika infection in the continental United States, with zero confirmed cases of autochthonous transmission.^23^ As of February 17, 2016, nine confirmed and ten suspected cases of Zika infection were reported among pregnant women at the national level.^24^ To-date, zero and 3 confirmed Zika infections have been reported among pregnant women in Texas^24^ and Florida^26^ compared to 0.95 and 1.51 infections projected by the model. We expect reported cases of Zika infection among pregnant women to increase in the coming months as Zika continues to spread in Central and South America. While the model is not exhibiting precise validation, we propose there is validation of an assertion that Zika virus is unlikely to cause localized medical infrastructure strain in the continental United States.

There has been substantial media attention regarding Zika infection as a threat to pregnancy, prompted by WHO’s declaration of a Public Health Event of International Concern (PHEIC).^2^ Our simple impact assessment model is only partially validating at this early interval in the crisis, where Zika emergence is ongoing. This is likely due to lack of precision in population, birth rate, or dengue or Chikungunya infection statistics combined with greater social sensitivity to report Zika infection due to the threat to the unborn, a socially sensitive population. Despite data suggesting an association between primary Chikungunya infection and adverse perinatal outcomes^27^, we propose the general public in the United States is unaware of this association due to lack of media campaigns conducted by the Centers for Disease Control and Prevention. The emergence of Chikungunya in the western hemisphere was not declared a PHEIC by the World Health Organization. Therefore, the level of threat perception in the United States is different for Zika compared to Chikungunya. We propose this discrepancy in sensitivity to seek laboratory screening by pregnant women will likely play a role in signal enhancement in the months to come. Lastly, it is also possible the transmission dynamics of Zika emergence in the context of an immunologically naïve population differs from that observed thus far with Chikungunya emergence in Central and South America. The degree to which this simple model validates in the future may serve as an indicator of such difference in transmission kinetics.

### Conflict of Interest Disclosure

The authors have read the journal’s policy, and the authors of this manuscript have the following competing interests: JMW is an employee and equity holder in M2 Medical Intelligence, Inc. RWM is an employee and equity holder in RW Malone MD LLC.

### Financial Support

The authors gratefully acknowledge the University of Nevada-Reno for their support of this study. The funders had no role in study design, data collection and analysis, decision to publish, or preparation of the manuscript.

## Acknowledgments

The authors thank the reviewers for their consideration of this manuscript.

